# Functional interrogation of a SARS-CoV-2 host protein interactome identifies unique and shared coronavirus host factors

**DOI:** 10.1101/2020.09.11.291716

**Authors:** H.-Heinrich Hoffmann, William M. Schneider, Francisco J. Sánchez-Rivera, Joseph M. Luna, Alison W. Ashbrook, Yadira M. Soto-Feliciano, Andrew A. Leal, Jérémie Le Pen, Inna Ricardo-Lax, Eleftherios Michailidis, Yuan Hao, Ansgar F. Stenzel, Avery Peace, C. David Allis, Scott W. Lowe, Margaret R. MacDonald, John T. Poirier, Charles M. Rice

## Abstract

The ongoing SARS-CoV-2 pandemic has devastated the global economy and claimed nearly one million lives, presenting an urgent global health crisis. To identify host factors required for infection by SARS-CoV-2 and seasonal coronaviruses, we designed a focused high-coverage CRISPR-Cas9 library targeting 332 members of a recently published SARS-CoV-2 protein interactome. We leveraged the compact nature of this library to systematically screen four related coronaviruses (HCoV-229E, HCoV-NL63, HCoV-OC43 and SARS-CoV-2) at two physiologically relevant temperatures (33 °C and 37 °C), allowing us to probe this interactome at a much higher resolution relative to genome scale studies. This approach yielded several new insights, including unexpected virus and temperature specific differences in Rab GTPase requirements and GPI anchor biosynthesis, as well as identification of multiple pan-coronavirus factors involved in cholesterol homeostasis. This coronavirus essentiality catalog could inform ongoing drug development efforts aimed at intercepting and treating COVID-19, and help prepare for future coronavirus outbreaks.

**HIGHLIGHTS:** Focused CRISPR screens targeting host factors in the SARS-CoV-2 interactome were performed for SARS-CoV-2, HCoV-229E, HCoV-NL63, and HCoV-OC43 coronaviruses.

Focused interactome CRISPR screens achieve higher resolution compared to genome-wide screens, leading to the identification of critical factors missed by the latter.

Parallel CRISPR screens against multiple coronaviruses uncover host factors and pathways with pan-coronavirus and virus-specific functional roles.

The number of host proteins that interact with a viral bait protein is not proportional to the number of functional interactors.

Novel SARS-CoV-2 host factors are expressed in relevant cell types in the human airway.

## INTRODUCTION

The ongoing coronavirus disease 2019 (COVID-19) pandemic has claimed the lives of nearly one million people worldwide and remains uncontrolled in several countries, presenting an urgent global health crisis. Coronaviruses are positive-sense RNA viruses that infect a variety of vertebrate hosts. Seasonal coronaviruses endemic in humans cause mild, common cold-like symptoms. In recent years, however, novel coronavirus strains circulating in animal reservoirs have spilled over into the human population causing severe disease. Three notable coronavirus outbreaks in the 21^st^ century include SARS-CoV in 2003, MERS-CoV in 2012, and SARS-CoV-2 in late 2019. Although the spread of SARS-CoV and MERS-CoV was limited, SARS-CoV-2, the etiological agent of COVID-19, remains widespread. With three known spillover events in recent history, this virus family has proven its potential to cross species, spread rapidly and take numerous lives.

The urgency of the ongoing SARS-CoV-2 pandemic has advanced coronavirus research at unprecedented speed. Two groups performed genome-wide CRISPR-Cas9 screens to identify host factors required by SARS-CoV-2 (Heaton et al., 2020; Wei et al., 2020) with the hope that these screens would not only uncover the biology underlying this virus, but also identify druggable host factors that could be targeted to prevent virus spread or lessen disease. Other groups conducted drug repurposing screens using compound libraries enriched with FDA-approved or clinical phase drug candidates to identify off-the-shelf interventions (Dittmar et al., 2020; Zhou et al., 2020). A complementary strategy to these screens is to nominate candidate host factors that are predicted to interact with viral proteins through immunoprecipitation mass spectrometry (IP-MS) of affinity tagged viral proteins. Indeed, one of the earliest discovery-based SARS-CoV-2 reports to appear in press was a study that aimed to map the SARS-CoV-2 host protein interactome using this technique (Gordon et al., 2020). This dataset highlighted potential targets to explore for drug repurposing based on the hypothesis that host proteins that directly or indirectly bind SARS-CoV-2 proteins are more likely to be required to support infection or spread relative to the rest of the proteome. Similar protein interactome studies in different contexts and with varying degrees of overlap between datasets soon followed (Stukalov et al., 2020). While there is a pressing need to better understand SARS-CoV-2 biology and identify potential therapeutic targets, it is also critical to obtain and interpret this knowledge in the broader context of coronavirus biology to prepare for potential future outbreaks. We therefore aimed to use CRISPR-Cas9 technology to interrogate SARS-CoV-2 host factor requirements in a broader context that included additional members of the coronavirus family.

CRISPR-Cas9 forward genetics and data from IP-MS studies can be combined in a high-coverage hypothesis-driven sgRNA screening campaign (Kelly et al., 2020). Focused libraries offer several advantages: 1) increased coverage (smaller libraries allow interrogation of many sgRNAs per gene, thereby lowering the rate of false-negatives), 2) increased resolution (smaller libraries ensure higher screen representation at every point of the screen), and 3) a wider range of screening conditions (smaller libraries are more compatible with highly parallel testing of multiple technical and biological conditions) (DeWeirdt et al., 2020; Doench, 2018; Parnas et al., 2015).

In this study, we leveraged the power of high resolution pooled CRISPR-Cas9 screens by generating and screening a focused sgRNA library targeting 332 host proteins identified by Gordon et al. as high-confidence SARS-CoV-2 protein interactors (Gordon et al., 2020). This focused approach allowed us to interrogate 10 sgRNAs per gene, thereby achieving near-saturating coverage of candidate targets. Furthermore, the reduced library complexity allowed us to systematically expand our screening efforts across a panel of four related human coronaviruses representing both alpha-(HCoV-229E and HCoV-NL63) and beta-coronaviruses (HCoV-OC43 and SARS-CoV-2). We conducted these screens at 33 °C and 37 °C, consistent with the native environment of the upper and lower airway, respectively. This systematic approach allowed us to uncover host proteins that are required for infection by all four tested human coronaviruses. Several of these host factors may serve as suitable targets for developing pan-coronavirus therapeutics. Furthermore, we uncovered previously unappreciated differences in host factor requirements among human coronaviruses, which could be leveraged for designing strain-specific therapeutic strategies. Overall, our study highlights the power of high-coverage CRISPR-Cas9 screens for systematic identification of host factors required for coronavirus infection and serves as a resource for efforts aimed at developing novel strategies to treat COVID-19, seasonal coronavirus infections, and perhaps future pandemic strains.

## RESULTS

### A focused CRISPR-Cas9 screen functionally validates host protein members of a SARS-CoV-2 host protein interactome

In a recent study, 26 of the 29 known SARS-CoV-2 proteins were individually expressed in HEK293T cells and immunoprecipitated to identify interacting host proteins (Gordon et al., 2020). After stringent filtering, 332 high-confidence host protein interactors were nominated. Here, we designed a custom-made sgRNA library and performed a series of CRISPR-Cas9 forward genetic screens to determine which of these 332 host proteins may be required for SARS-CoV-2 infection and virus-induced cytopathogenicity.

We constructed a lentiviral transfer plasmid library encoding 3,944 sgRNAs designed to disrupt all 332 human genes identified in the abovementioned SARS-CoV-2 interactome (**Figure 1A**). The library includes 10 sgRNAs targeting each gene, 314 safe-targeting sgRNAs (which cut the genome at non-genic regions (Morgens et al., 2017)), and 310 sgRNAs targeting genes essential for cell survival or proliferation (Hart et al., 2017; Michlits et al., 2020). We used this library to perform a pooled CRISPR-Cas9 gene disruption screen in Huh-7.5 hepatoma cells stably expressing Cas9 (Huh-7.5-Cas9) (**Figure 1B**). Huh-7.5 cells were chosen because they express the SARS-CoV-2 cellular receptor, angiotensin-converting enzyme 2 (ACE2), as well as transmembrane serine protease 2 (TMPRSS2), which is involved in SARS-CoV-2 entry (**Figure S1A**) (Hoffmann et al., 2020). Furthermore, these cells express 329/332 (99%) of the host proteins in the SARS-CoV-2 interactome at levels similar to highly susceptible Calu-3 cells (**Figure S1A)** (Wyler et al., 2020). Importantly, we confirmed that Huh-7.5 cells were readily infected and displayed virus-induced cytopathic effect (CPE) by multiple coronaviruses, including SARS-CoV-2, thereby validating these cells as a robust system for studying members of the *Coronaviridae* family (**Figure S1B**).

**Figure 1.**
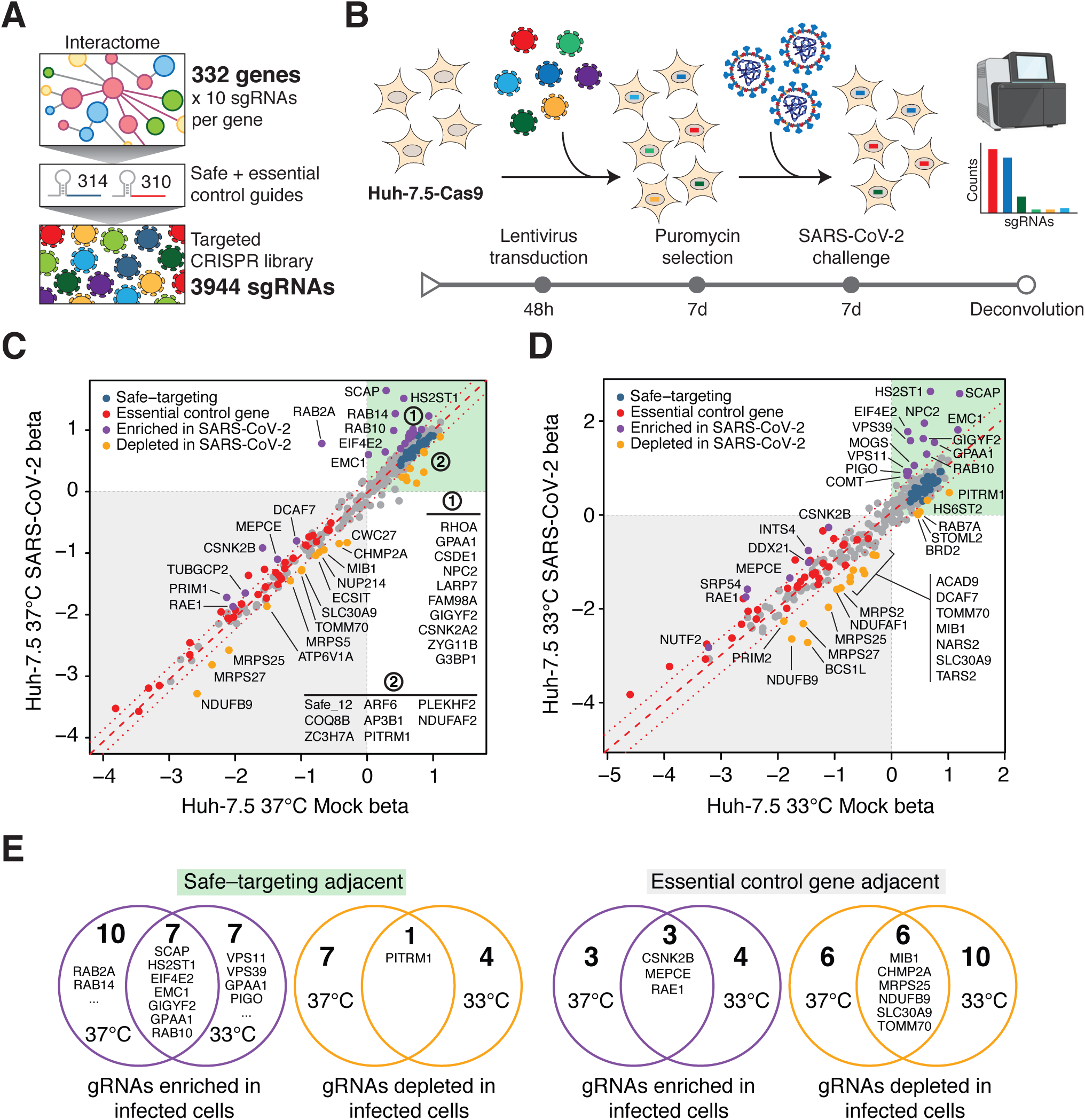
A SARS-CoV-2 protein interactome CRISPR screen identifies functional SARS-CoV-2 protein interactors. **(A)** An sgRNA library targeting 332 interactome genes along with 314 safe and 310 essential control sgRNAs was used to create a targeted lentiviral library. **(B)** CRISPR screening workflow. Cas9 expressing cells are transduced with lentivirus encoding the sgRNA library and subsequently selected with SARS-CoV-2. Surviving cells along with mock controls are subject to sgRNA sequencing. **(C)** Scatterplot of gene essentiality scores for a SARS-CoV-2 interactome screen performed in Huh-7.5 cells cultured at 37 °C. **(D)** Scatterplot of a SARS-CoV-2 interactome screen performed in Huh-7.5 cells at 33 °C. **(E)** Venn diagrams depicting overlap for enriched or depleted sgRNAs at 37 °C and 33 °C. Genes with beta scores similar to safe-targeting sgRNAs (left) or essential genes (right) are indicated.

Next, we performed a series of genetic screens by transducing Huh-7.5-Cas9 cells with the SARS-CoV-2 interactome sgRNA library followed by puromycin selection and expansion for one week prior to SARS-CoV-2 challenge. To identify potential temperature-dependent differences in host factor requirements, SARS-CoV-2 selection screens were performed at two temperatures, 33 °C and 37 °C, to mimic the temperatures of the upper and lower airway, respectively (V’kovski et al., 2020). Library-transduced cells that survived SARS-CoV-2 infection were harvested after seven (33 °C) and eleven (37 °C) days followed by genomic DNA isolation and screen deconvolution via next generation sequencing (Methods). In this context, cells transduced with sgRNAs targeting host factors required for viral infection or virus-induced CPE should survive and enrich within the infected population while cells expressing safe-targeting sgRNAs or sgRNAs targeting genes inconsequential to virus infection are expected to die and deplete over time. Similarly, cells transduced with sgRNAs targeting genes essential for cell survival should deplete independently of SARS-CoV-2 infection.

The overall technical performance of these genetic screens was excellent (**Figure S1C-F**). We confirmed that 3,921/3,944 (99.4%) of sgRNAs were recovered from the plasmid library and that all screen libraries were sequenced to saturation (**Figure S1C)**. Correlation analyses demonstrated that biological replicates for each condition clustered together and shared a high correlation coefficient (≥ .97), as expected (**Figure S1D)**.

Receiver operating characteristic (ROC) curves, which measure the technical performance of genetic screens based on the degree to which the screen can distinguish between negative control (safe-targeting) and positive control (essential gene-targeting) sgRNAs, were excellent (**Figure S1E-F**). Of note, incorporating safe-targeting and essential gene-targeting sgRNAs allows for better internal calibration and effect size estimation of CRISPR-based genetic screens (Methods). Specifically, we used these built-in controls to establish a gene essentiality (beta) score (Li et al., 2015), which allowed us to first stratify interactome targets based on their effects on cellular fitness under “mock” (uninfected) conditions followed by identifying high-confidence gene “hits” in infected cells (Methods). Genes with beta scores similar to essential genes (which influence cell survival with or without infection) may be confounded by effects on cellular fitness whereas genes with beta scores similar to safe-targeting sgRNAs (which only influence survival during viral infection) are more likely to be true positives. Using these parameters, we identified 36 and 43 genes that significantly influenced SARS-CoV-2-induced cell death at 33 °C and 37 °C, respectively (**Figure 1C-D and Table S1C, E**). Of these, 21 (33 °C) and 23 (37 °C) genes scored as potential host factors that promote SARS-CoV-2-induced cell death (purple dots in **Figure 1C-D**), while 21 (33 °C) and 20 (37 °C) genes scored as putative antagonists of the virus life cycle (orange dots in **Figure 1C-D**). In contrast, safe-targeting and essential gene-targeting sgRNAs behaved similarly across mock and SARS-CoV-2 conditions (blue and red dots in **Figure 1C-D**, respectively). These results suggest that the SARS-CoV-2 host protein interactome is composed of functionally relevant factors that play critical roles during the life cycle of this virus.

As shown in **Figures 1C and 1D**, we noted significant enrichment in sgRNAs targeting sterol regulatory element-binding protein (SREBP) cleavage-activating protein (SCAP), heparan sulfate 2-O-sulfotransferase 1 (HS2ST1), eukaryotic translation initiation factor 4E type 2 (EIF4E2), the Rab family of small GTPases (e.g., RAB2A, RAB10, and RAB14), members of the homotypic fusion and vacuole protein sorting (HOPS) tethering complex (e.g., VPS39 and VPS11), and ER Membrane Protein Complex Subunit 1 (EMC1). Additional significantly enriched genes are highlighted in purple in **Figures 1C and 1D**. These results indicate that these SARS-CoV-2 protein interactors are essential for virus infection or virus-induced cell death. In contrast, we observed negative enrichment in mammalian mitochondrial ribosomal small subunit (MRPS) genes (e.g., MRPS2, MRPS5, MRPS25, and MRPS27), suggesting that loss of these genes sensitize cells to direct or indirect virus-induced cell death. Additional significantly depleted genes are highlighted in orange in **Figures 1C and 1D**.

We also observed hits that were common to screens performed at either 33 °C or 37 °C (**Figure 1E**), as well as hits that were specific to one of the two temperatures. An intriguing outcome is the apparent differential requirement for two Rab GTPases, RAB2A and RAB14, which appear to be essential for SARS-CoV-2 infection and virus-induced CPE at 37 °C but not at 33 °C. In contrast, members of the HOPS complex (VPS11 and VPS39) and proteins involved in glycosylphosphatidylinositol (GPI) anchor biosynthesis (PIGO and GPAA1) appear to be required at 33 °C but not at 37 °C (**Figure 1E**). These findings may reflect differences in the SARS-CoV-2 entry pathway at 33 °C and 37 °C.

Lastly, we found that positively selected hits were typically enriched in the region of safe-targeting sgRNAs, while negatively selected hits tended to be enriched in the region of control genes with deleterious effects (**Figure 1E**).

In conclusion, our SARS-CoV-2 interactome screen identified a compendium of host genes that can be disrupted without affecting cell viability in the absence of infection, yet seem to play essential roles during SARS-CoV-2 induced CPE (**Figure 1C-E**). Such genes could represent excellent candidates for the development of anti-SARS-CoV-2 therapeutics.

### Multiple coronaviruses require both obligate and likely dispensable SARS-CoV-2 host factors for infection

Viruses within the same family can have both shared and specific host factor requirements. Functionally distinguishing these requirements is critical to gain a complete understanding of virus-host biology and, on a practical level, can help identify targets for pan-viral therapies. Towards these goals, we expanded our SARS-CoV-2 interactome CRISPR-Cas9 screening campaign by performing complementary screens using three additional human coronaviruses: HCoV-229E, HCoV-NL63, and HCoV-OC43. To visualize the degree of enrichment in genes that scored in the SARS-CoV-2 screens described above, we plotted the cumulative distribution of absolute z-scores for all five screens (**Figure 2A**). Each of these screens were significantly enriched in the subset of genes that scored in the initial SARS-CoV-2 screens as compared to the complete library.

**Figure 2.**
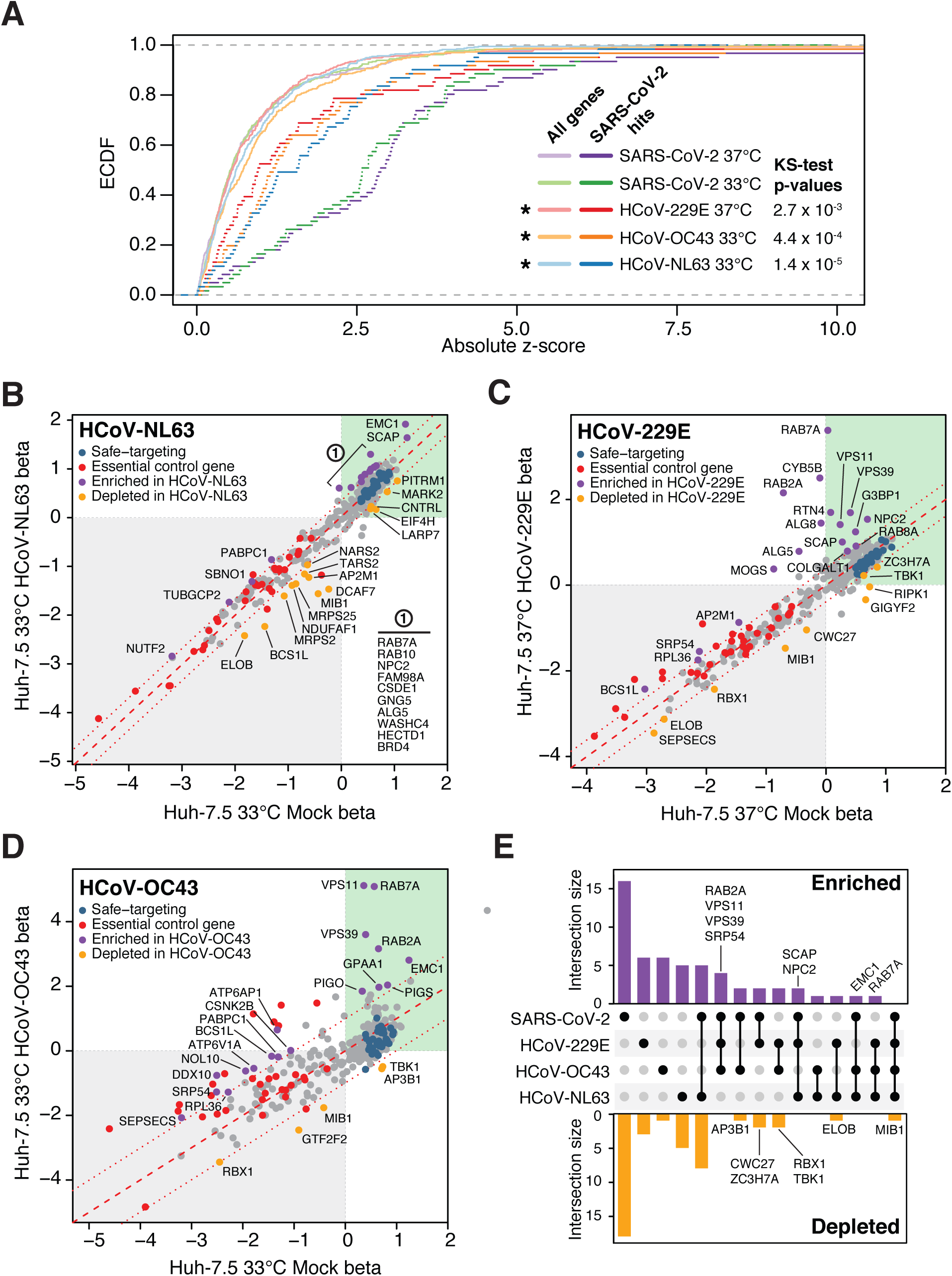
Functional interrogation of a SARS-CoV-2 protein interactome with additional coronaviruses identifies pan-coronavirus and coronavirus-specific host factors. **(A)** Empirical cumulative distribution function (ecdf) plot of absolute z-scores for SARS-CoV-2 hits across screens for the four coronaviruses as indicated (dark color) compared to all tested genes (All genes, light color). Statistical significance of enrichment determined by two-sided Kolmogorov–Smirnov (K-S) test. **(B-D)** Scatterplot of gene essentiality scores for a SARS-CoV-2 protein interactome screened in Huh-7.5 cells with **(B)** HCoV-NL63 at 33 °C, **(C)** HCoV-229E at 37 °C and **(D)** HCoV-OC43 at 33 °C. **(E)** UpSet plot showcasing hits overlapping in screens across all four viruses. SARS-CoV-2 hits are inclusive of both temperatures, 33 °C and 37 °C. Select genes for enriched or depleted sgRNAs are indicated.

These screens also identified Rab GTPases, components of the HOPS complex, and EMC1 as host factors required for coronavirus infection and virus-induced CPE (**Figure 2B-D**). In addition to hits common among different coronaviruses, several strain-specific hits were also identified (**Figure 2E**). For example, HCoV-OC43, for which there is currently no protein receptor described and entry has been associated with binding to 9-O-acetylated sialic acids (Hulswit et al., 2019), displayed a unique requirement for GPI anchor biosynthesis as evidenced by the fact that GPAA1, PIGO, and PIGS scored as significant hits in the screen (**Figure 2D and Figure S4A**).

In addition to sgRNAs that enrich in virus-infected cell populations (suggesting the genes targeted by these sgRNAs support virus infection), we also identified sgRNAs that were depleted in virus-infected cultures (suggesting the genes targeted by these sgRNAs could have an antiviral function). For example, Mindbomb 1 (MIB1), an E3 ubiquitin-protein ligase, was specifically depleted in the context of infection with all four viruses. Although MIB1 disruption was deleterious in both mock infected and virus infected cells, MIB1 disruption was consistently more toxic in the context of virus infection. MIB1 is known to play a role in ‘Lys-63’-linked polyubiquitination of TBK1 kinase, which is thought to mediate its activation (Li et al., 2011). Consistent with a potential role in this pathway, sgRNAs targeting TBK1 were negatively enriched in HCoV-229E and HCoV-OC43 screens (**Figure 2C-D**). This might suggest a role for MIB1 and TBK1, a signal integrator of multiple RIG-like receptors (RLRs) and positive regulator of IRF3, in establishing an antiviral state that controls coronavirus infection. This could explain why, in the absence of MIB1 and TBK1, cells are more susceptible to virus-induced CPE.

### Functional interactors are not proportionally distributed between bait proteins

We next asked how the functional hits we identified related to the viral bait proteins with which they interact. To visualize functional and network relationships, we overlaid our CRISPR screening results from all viral screens in this study onto a bait-centric interactome map (**Figure 3A**). The distribution of functional hits across different bait proteins was apparently non-random; instead, we observed that certain bait proteins appeared to be enriched in functional hits.

**Figure 3.**
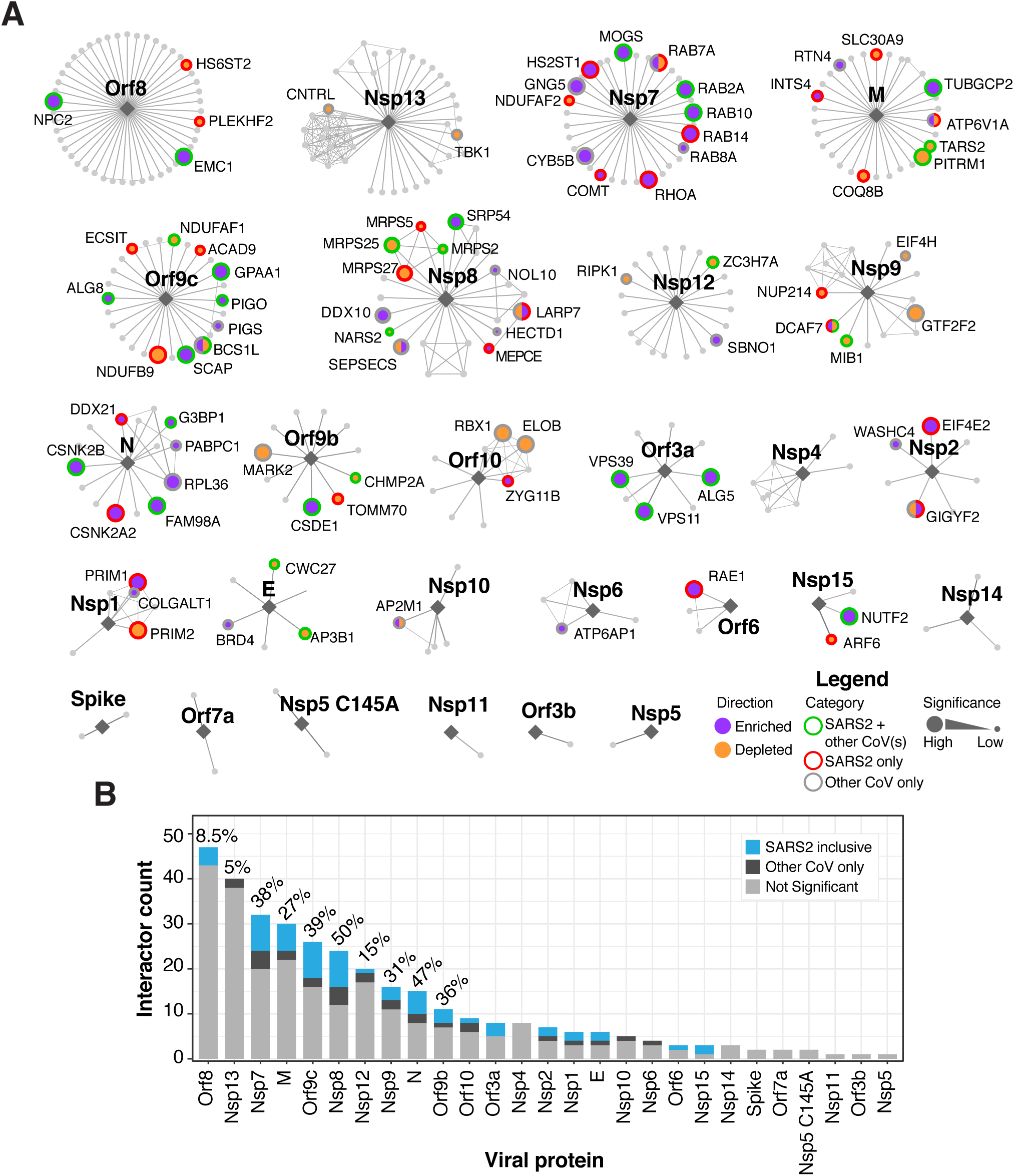
Network overlay of interactome CRISPR screening. **(A)** Interactome network organized by virus bait protein from largest to smallest number of putative interactors. Enriched or depleted hits are highlighted in labeled purple or orange filled circles, respectively. Border colors denote SARS-CoV-2 hits that are shared with any other coronavirus screen (green), hits exclusive to SARS-CoV2 (red) or hits exclusive to other HCoVs (gray) with size of node scaled with significance. **(B)** Barplot of nodes per viral protein with all SARS-CoV-2 shared and exclusive hits indicated in blue. Hits for other CoVs are shown in dark gray. Percentage of genes that scored as a hit in any screen are indicated for all genes with greater than 10 edges.

We subsequently explored whether the number of host protein interactions for a given bait protein was predictive of the percentage of functional interactors (**Figure 3B**). There was no evident correlation between the number of predicted host protein binding partners of a given viral protein and the number of functional interactors. We found that some viral baits that interact with many host proteins (e.g., Orf8 and Nsp13) have relatively few functional interactors, whereas others (e.g., Nsp7, Nsp8, and Orf9c) have many. Overall, most of the functional hits identified in our pan-coronavirus screens seem to be required for SARS-CoV-2 infection. This was expected given that our screens were based on a SARS-CoV-2 protein interactome. We were surprised, however, to find that there are SARS-CoV-2 protein interactors that are essential for one or more of the other coronaviruses but *not* essential for SARS-CoV-2 (e.g., DDX10, RPL36, GNG5, and CYB5B; **Figure 2E, 3A**). Altogether, these data suggest that the relationship between physical interactome screens and functional genetic screens is not strictly concordant. Some viral proteins may interact with redundant factors whose depletion is compensated. However, 27% of SARS-CoV-2 interacting proteins functionally influenced the outcome of infection, suggesting that this high-confidence interactome is substantially enriched in functional host factors.

## DISCUSSION

Host:pathogen protein:protein interaction (PPI) networks derived from immunoprecipitation mass spectrometry (IP-MS) data have proven valuable for illustrating the complex nexus between viral and cellular proteins (Gordon et al., 2020; Sadegh et al., 2020; Stukalov et al., 2020). PPI networks can predict hundreds of potential interactions; however, secondary validation of each interaction is often a labor-intensive process. Moreover, it is not possible to assign functional significance to any particular *bona fide* interaction based on co-purification or co-immunoprecipitation alone in part due to possible artifacts caused by overexpression outside of the viral context. Functional experiments are therefore needed to assess the biological significance of each host protein node in the network.

We probed a SARS-CoV-2 PPI network consisting of 332 high-confidence host factors by systematically screening multiple coronaviruses using a novel, high-density, pooled sgRNA library. First, we interrogated the functional role of each predicted member of the network in the presence or absence of SARS-CoV-2 infection, which allowed us to identify host factors that appear to be dispensable for Huh-7.5 proliferation but play a functional role in SARS-CoV-2 infection. Of the predicted interactome, we were able to assign a functional role for 87/332 (26%) unique genes. This finding is in contrast to two recent preprint studies that performed genome-scale CRISPR-Cas9 screens in VeroE6 and ACE2-reconstituted A549 cells, which failed to identify significant enrichment of the same PPI network (Heaton et al., 2020; Wei et al., 2020). We speculate that this difference may be due in part to well-established advantages of focused screens, including 1) improved signal-to-noise ratios in the absence of sgRNAs targeting genes that are likely to dominate the screen (e.g., the SARS-CoV-2 cellular receptor, ACE2), 2) increased coverage by including more sgRNAs per gene (ten vs. four), 3) feasibility of exploring multiple screening conditions (e.g., different viruses and temperatures), and 4) higher screen representation (i.e., > 1,500-fold vs. < 750-fold) ((DeWeirdt et al., 2020; Doench, 2018; Parnas et al., 2015)). Thus, by deeply probing individual interactome proteins using a focused sgRNA library coupled with functional readouts across multiple conditions allowed us to identify many host factors required for SARS-CoV-2 infection that were missed in genome-scale CRISPR screens (Heaton et al., 2020; Wei et al., 2020).

While host proteins associated with specific viral bait proteins (e.g., Nsp7, Nsp8, and Orf9c) appear to be enriched in functionally required host factors, we note that the greatest degree of co-essentiality was predominantly observed between functionally-related host factors instead of between factors associated with the same viral bait protein (e.g., SCAP and NPC2, both of which are involved in cholesterol homeostasis but interact with Orf9c and Orf8, respectively). This emphasizes the complementary role played by IP-MS data and functional genomics in identifying viral proteins that serve as hubs for multiple host factors that may participate in diverse molecular processes.

In addition to the pandemic SARS-CoV-2 strain, we performed parallel screens for three seasonal human coronaviruses (HCoV-NL63, HCoV-229E and HCoV-OC43), which allowed us to compare and contrast virus-specific host factor requirements and identify potential pan-coronavirus host factors. Many genes from the SARS-CoV-2 PPI network were essential across multiple coronaviruses tested (40/332; 12%), suggesting functional conservation across host:coronavirus PPIs. Surprisingly, though, we found that some predicted SARS-CoV-2 interactors appeared *not* to play a role in SARS-CoV-2 infection but were essential for other coronaviruses (**Figure 4A**). This may be due to promiscuity in protein-protein interactions, evolutionary divergence in the functional requirement for a given host factor, redundant factors (e.g., SARS-CoV-2 might rely on redundant factors for specific viral life cycle steps while other coronaviruses might rely on a single factor), or the strict threshold—the ability to survive infection—that must be crossed to score as a hit in these screens. For example, it is possible that inactivation of specific host factors impairs SARS-CoV-2 infection to some extent but the virus nevertheless kills the cell.

**Figure 4.**
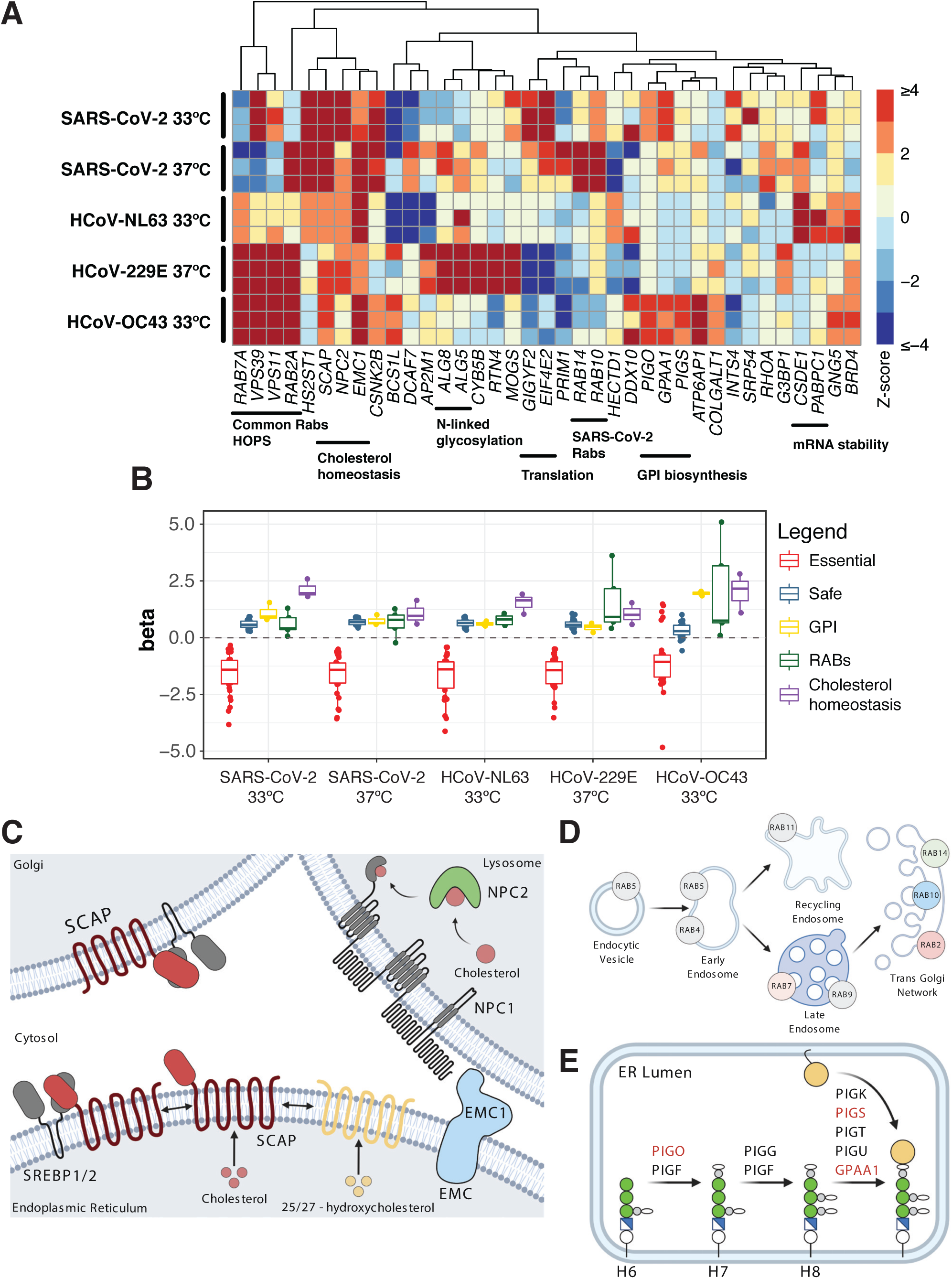
Pathway focused views of high confidence CRISPR hits. **(A)** Heatmap of z-scores for the union of positively-enriched coronavirus host factors across experimental replicates in all infection conditions. Functionally-related genes are highlighted. **(B)** Distribution of gene essentiality (beta) scores for the pathways highlighted in panel A. **(C)** SCAP/SREBP pathway regulating cholesterol homeostasis. **(D)** Rab GTPases and intracellular vesicle trafficking. **(E)** Biosynthesis of GPI-anchored proteins. Diagrams created with BioRender.com. Modified from Maeda et al., 2017, Shimano and Sato, 2017, and Stenmark, 2009.

To illustrate the power and versatility of our screening approach, we discuss below a specific example of pan-coronavirus and strain-specific host factors uncovered in our studies. The rest of the screening data can be found in **Table S1**.

### SCAP is a required and druggable SARS-CoV-2 and pan-coronavirus host factor

A central goal of this study was to use a predicted SARS-CoV-2 interactome to identify host factors that are required for virus infection or virus-induced CPE by multiple coronaviruses. We identified SCAP as a pan-coronavirus host factor required for all four of the coronaviruses tested in this study (**Figure 4A-C**). SCAP regulates lipid and cholesterol homeostasis by sequestering SREBPs in the ER in the presence of sterols, thus preventing their transport to the Golgi apparatus in COPII-coated vesicles where SREBPs are normally activated by proteolytic cleavage and subsequently shuttled to the nucleus to regulate gene expression (Luo et al., 2020). It is possible that multiple coronaviruses hijack this transport pathway as part of their life cycle or that coronavirus infection is not compatible with cell membrane properties in a low cholesterol state. Interestingly, Niemann–Pick intracellular cholesterol transporter 2 (NPC2) and EMC1 were also enriched across multiple coronavirus screens (**Figure 4A-C**). NPC2 normally plays a role in trafficking cholesterol out of lysosomes (Infante et al., 2008), and disruption of NPC2 could decrease cholesterol availability, which would prevent sequestration of SREBPs by SCAP and thus, phenocopy SCAP loss of function. EMC1 has been implicated in the exit of simian virus 40 (SV40) from the ER (Bagchi et al., 2016), in the infection of many flaviviruses (Marceau et al., 2016; Savidis et al., 2016; Zhang et al., 2016), and as part of the EMC complex in cholesterol biosynthesis (Volkmar et al., 2019). SCAP and NPC2 have not been previously identified in other host:pathogen forward genetic screens; however, an upstream cholesterol transporter, NPC1, was identified as an essential host factor for Ebola virus (Carette et al., 2011). The SCAP:SREBP complex is further regulated by oxysterol sensors, including insulin induced gene 2 (INSIG2), which sequesters SCAP:SREBP in the presence of oxysterols like 25-hydroxycholesterol (25OHC) (Radhakrishnan et al., 2007). It is intriguing to speculate that oxysterols or direct SCAP inhibitors, such as fatostatin (Kamisuki et al., 2009) or its derivatives, could have antiviral properties. Consistent with this possibility, one study showed that 27-hydroxycholesterol (27OHC) exerted SARS-CoV-2 antiviral activity in VeroE6 cells at 10 μM, and that overall oxysterols were lower in patients with moderate and severe COVID-19 symptoms compared to controls (Marcello et al., 2020). Another study showed that 25OHC had potent SARS-CoV-2 antiviral activity at similar concentrations (Zang et al., 2020). These findings are in strong agreement with work from multiple groups that has demonstrated that 25OHC is a potent inhibitor of many enveloped viruses (Liu et al., 2013). Importantly, these regulators of lipid and cholesterol homeostasis are all constitutively expressed in Huh-7.5 cells (**Figure 4B**), as well as in cells and tissues known to be infected by SARS-CoV-2 (**Figure S2**). Collectively, these results nominate regulators of lipid and cholesterol homeostasis as potential critical regulators of coronavirus infection and further suggest that direct targeting of SCAP may be a potential pan-coronavirus therapeutic strategy.

### Differential requirement of Rab GTPases among members of the *Coronaviridae* family

Another central goal of this study was to probe the SARS-CoV-2 interactome across multiple coronaviruses to identify host factors required for virus infection or virus-induced CPE by specific strains (e.g., SARS-CoV-2-specific factors that are dispensable for HCoV-NL63, HCoV-229E, and/or HCoV-OC43 infection). By comparing two temperatures and four coronaviruses in parallel, we uncovered striking heterogeneity in the requirement for different Rab GTPases, which varied by coronavirus and temperature (**Figure 4A-B, D**). Specifically, we identified a Rab GTPase module consisting of RAB2A and RAB7A that appears to be critical for infection and virus-induced CPE by HCoV-229E, HCoV-OC43, and HCoV-NL63, but that seems to be partially dispensable for SARS-CoV-2 infection, at least at lower temperatures (**Figure 4A**). In accordance with these results, we identified the RAB10 and RAB14 GTPases as critical SARS-CoV-2-specific host factors while RAB5C appears to be specific for HCoV-OC43 infection (**Table S1**). This differential requirement is not due to differences in basal expression levels of Rab GTPases, as mRNAs for all five Rab proteins are uniformly expressed in Huh-7.5 cells, as well as in cells and tissues known to be infected by SARS-CoV-2 (**Figure 4B, Figure S2**). These results suggest that different coronaviruses may hijack specific Rab GTPase modules in host cells to carry out their infectious cycle, raising the possibility that targeting these modules could have therapeutic benefits. Furthermore, given the central role that Rab GTPases play in orchestrating complex intracellular trafficking, the observation that SARS-CoV-2 and HCoV-NL63, which both utilize ACE2 for cellular entry (Hoffmann et al., 2020; Hofmann et al., 2005), have different requirements for Rab GTPases suggests that even coronaviruses that engage the same cellular receptor may differ in their entry pathways in a context-dependent manner. Moreover, the temperature-specific differences we observed among the same virus (SARS-CoV-2) raises the intriguing hypothesis that there may be differences in the infection process across the temperature gradient spanning the upper and lower airway. Recent studies have nominated additional SARS-CoV-2 entry factors, including (Gu et al., 2020). One potential explanation for the temperature-specific differences we observed in entry pathways could be differential use of entry factors at 33 °C and 37 °C. In addition to virus- and temperature-specific differences in the requirement of Rab GTPases, we found that disruption of GPI anchor biosynthesis uniquely affected HCoV-OC43 (**Figure 4A-B, E**), raising the tantalizing possibility that HCoV-OC43 engages one or more GPI-anchored host factors, potential receptors, or co-receptors, that facilitate entry upon binding to 9-O-acetylated sialic acids (Hulswit et al., 2019).

This study leverages the power of CRISPR-Cas9 forward genetic screens to functionally probe a high-confidence SARS-CoV-2 host protein interactome across multiple coronaviruses and temperatures. In doing so, our study delineates the complex functional interplay between the interacting viral and host factors defined by Gordon et al. (Gordon et al., 2020) and nominates a set of core biological programs that play critical roles during infection by SARS-CoV-2 and other coronaviruses.. We found that a significant fraction of putative interactors in the SARS-CoV-2 interactome are indeed required for viral infection and virus-induced CPE by the pandemic SARS-CoV-2 and other endemic coronaviruses (HCoV-NL63, HCoV-229E, and HCoV-OC43). Interestingly, we identified one class of host factors that appear to be critical for infection by all coronaviruses tested (e.g., regulators of cholesterol homeostasis) and other classes of factors that seem to be required only by specific coronaviruses (e.g., Rab GTPases and GPI-anchoring proteins). These findings fill an important knowledge gap between predicted and functional virus-host interactomes and may help to focus attention on the genes and pathways in the SARS-CoV-2 interactome that appear to have the greatest functional significance during early infection steps. More broadly, this study presents a generalizable approach to interrogate host:pathogen PPI networks with functional genomics. We predict that the compendium of host factors and pathways described in this work will provide novel insights into the infection mechanics of SARS-CoV-2 and seasonal coronaviruses, while simultaneously nominating candidate targets for antiviral strategies.

## LIMITATIONS

While our study sheds new light on how host proteins are co-opted during coronavirus infection, it is not without limitations. As with all pooled CRISPR gene disruption screens, we expect that functionally redundant genes may not score (Ewen-Campen et al., 2017)). Additionally, the results of these genetic screens should not be interpreted to validate specific physical associations, as it is a purely functional readout. Additional biochemical studies would be necessary to test each predicted interaction. Nevertheless, we identified robust pan-coronavirus and discrete strain-specific host factors that scored with up to 10 sgRNAs scoring per gene, validating the utility of this near-saturation CRISPR-based genetic screening approach. We acknowledge that Huh-7.5 cells, which were chosen based on their unique capacity to support infection by multiple coronaviruses and also because they constitutively express 324/332 of the SARS-CoV-2 interactome factors, are different than the HEK293T system used by Gordon et al. to generate the PPI network probed in this study and are not airway cells. However, this robust and experimentally tractable system offers a unique advantage to directly compare hard-wired host factor requirements across multiple viruses in a cell-type- and tissue-type-agnostic manner, which can then be validated using gold-standard primary cells or animal models. Future studies will be required to dissect the specific requirement for each member of the PPI network validated in this study using additional models of coronavirus infection. Another potential limitation of our study is that the screening strategy is not optimal for assessing the functionality of host factors beyond their role in modulating cell survival. In other words, our current experimental set up does not allow for probing the functionality of interacting proteins or otherwise critical host factors that act in late stages of the viral life cycle. Furthermore, our approach cannot identify genes that play important roles in immune modulation and pathogenesis. Lastly, our genetic screens were not genome-wide and, consequently, should not be interpreted as a comprehensive catalog of potential coronavirus host factors. Future studies that employ focused CRISPR-Cas9 libraries targeting other pathways of interest and/or high coverage genome-wide libraries will undoubtedly extend this catalog of host factors hijacked by human coronaviruses.

## ACKNOWLEDGEMENTS

This work was initiated and conducted under unusual circumstances. As New York City and much of the world was sheltering in place to reduce the spread of SARS-CoV-2, all of the authors here were sustained during the shutdown by generous funding intended for related and unrelated work. During this time we were fortunate to obtain funding from government and charitable agencies that allowed this COVID-19 work to continue. For funding directly related to these COVID-19 efforts, we thank the G. Harold and Leila Y. Mathers Charitable Foundation and the Bawd Foundation for their generous awards. We also thank Fast Grants (www.fastgrants.org), a part of Emergent Ventures at the Mercatus Center, George Mason University. Research reported in this publication was supported in part by the National Institute of Allergy and Infectious Diseases of the National Institutes of Health under Award Number R01AI091707. The content is solely the responsibility of the authors and does not necessarily represent the official views of the National Institutes of Health. We further received funding for COVID-19 related work from an administrative supplement to U19AI111825. As mentioned above, authors were also supported with non-COVID-19 funding by the following awards, foundations, and charitable trusts: P01CA196539, R01CA190261, R01CA204639, R01CA21344, U01CA213359, R01AI143295, R01AI150275, R01AI143295, R01AI116943, P01AI138938, P30CA008748, P30CA016087, R03AI141855, R21AI142010, W81XWH1910409, EMBO Fellowship ALTF 380-2018, F32AI133910, the Robertson Foundation, the Center for Basic and Translational Research on Disorders of the Digestive System through the generosity of the Leona M. and Harry B. Helmsley Charitable Trust, the Leukemia and Lymphoma Society (LLS-SCOR 7006-13), the Rockefeller University and St Jude Children’s Research Hospital Collaborative on Chromatin Regulation in Pediatric Cancer (A11576), and an Agilent Technologies Thought Leader Award. S.W.L. is the Geoffrey Beene Chair of Cancer Biology and a Howard Hughes Medical Institute Investigator. Y.M.S.F was supported by the Damon Runyon-Sohn Pediatric Cancer Fellowship (DRSG-21-17). F.J.S-R is a HHMI Hanna Gray Fellow and was partially supported by an MSKCC Translational Research Oncology Training Fellowship (NIH T32-CA160001). We also thank the NYU Langone Health Genome Technology Center. We also wish to thank Laura Whitman, David Weiss, Joseph Marino, and Leo Brizuela from Agilent Technologies for rapid high-fidelity oligo synthesis support during the peak of the COVID-19 pandemic, Johannes Zuber (Research Institute of Molecular Pathology, Vienna) for support with VBC sgRNA scores, and Aileen O’Connell, Santa Maria Pecoraro Di Vittorio, Glen Santiago, Mary Ellen Castillo, Arnella Webson and Sonia Shirley for outstanding administrative or technical support.

## AUTHOR CONTRIBUTIONS

Conceptualization: HHH, WMS, FJSR, JTP, CMR

Methodology: HHH, WMS, FJSR, YMSF, JML, AL, JTP

Formal analysis: JML, YH, JTP

Investigation: HHH, WMS, AL, JLP, IRL, EM, AFS, JTP

Resources: CDA, SWL, JTP, CMR

Data curation: JML, JLP, JTP

Supervision: JTP, CMR

Visualization: FSR, JML, JTP

Writing – original draft: HHH, WMS, FJSR, JML, JTP

Writing – review & editing: HHH, WMS, FJSR, JML, JLP, IRL, MRM, JTP, CMR

Project Administration: AWA, AP, MRM

Funding acquisition: SWL, JTP, CMR

## SUPPLEMENTARY TABLES

**Table S1. CRISPR screen results**.

Table S1A: Huh-7.5 33 °C HCoV-NL63 meta-zscore.

Table S1B: Huh-7.5 33 °C HCoV-OC43 meta-zscore.

Table S1C: Huh-7.5 33 °C SARS-CoV-2 meta-zscore.

Table S1D: Huh-7.5 37 °C HCoV-229E meta-zscore.

Table S1E: Huh-7.5 37 °C SARS-CoV-2 meta-zscore.

Table S1F: mle plasmid gene summary.

Table S1G: mle plasmid sgrna summary.

**Table S2. Sequences of sgRNAs and primers**.

Table S2A: Gordon et al. SARS-CoV-2 Human Interactome CRISPR Library (sgRNA sequences).

Table S2B: Gordon et al. SARS-CoV-2 Human Interactome CRISPR Library cloning primers.

Table S2C: Primers used to prepare Illumina-compatible libraries for high-throughput sequencing of CRISPR screens.

Table S2D: Full sequence of sgRNA expression lentiviral vectors constructed and used in this study.

## LEAD CONTACT AND MATERIALS AVAILABILITY

Further information and requests for resources and reagents should be directed to and will be fulfilled by the Lead Contacts, John T. Poirier and Charles M. Rice.

## MATERIALS AND METHODS (Supplementary Tables here and here)

### Plasmids and sgRNA cloning

To generate stable Cas9-expressing cell lines, we used lentiCas9-Blast (Addgene, cat. #52962). To express sgRNAs, we used lentiGuidePurov2, a variant of lentiGuide-Puro (Addgene, cat. #52963) that contains an improved sgRNA scaffold based on Chen et al. 2013 (Chen et al., 2013). For sgRNA cloning, lentiGuidePurov2 was linearized with BsmBI (NEB) and ligated with BsmBI-compatible annealed and phosphorylated oligos encoding sgRNAs using high concentration T4 DNA ligase (NEB). All sgRNA sequences used are listed in Table S1. HIV-1 Gag-Pol and VSV-G plasmid sequences are available upon request.

### Cloning of the Gordon et al. SARS-CoV-2 Human Interactome CRISPR Library

sgRNA sequences (10 per gene) targeting 332 host factor proteins recently shown to interact with SARS-CoV-2 proteins (Gordon et al., 2020) (Table S1A) were designed using a combination of the Vienna Bioactivity CRISPR score (Michlits et al., 2020) and the Broad Institute sgRNA Designer tool (Doench et al., 2016). In addition, a subset of sgRNAs were hand-picked from the Wang-Sabatini genome-wide libraries (Wang et al., 2015). We also included 314 safe-targeting sgRNAs (Morgens et al., 2017) and 310 sgRNAs targeting essential genes (Hart et al., 2017; Michlits et al., 2020) for a total of 3,944 sgRNAs. We refer to this library as the ‘Gordon et al. SARS-CoV-2 Human Interactome CRISPR Library’. Oligo pools were synthesized by Agilent Technologies. The Gordon et al. SARS-CoV-2 Human Interactome CRISPR Library was cloned into lentiGuidePurov2 using a modified version of the protocol published by Doench et al (Doench et al., 2016) to ensure a library representation of > 10,000-fold. Briefly, the oligo pool was selectively amplified using barcoded forward and reverse primers that append cloning adapters at the 5’ and 3’ ends of the sgRNA insert (Table S1B), purified using the QIAquick PCR Purification Kit (Qiagen), and ligated into BsmBI-digested and dephosphorylated lentiGuidePurov2 using high-concentration T4 DNA ligase (NEB). A total of 4.8 μg of ligated lentiGuidePurov2 Gordon et al. SARS-CoV-2 Human Interactome CRISPR Library plasmid DNA was electroporated into Endura electrocompetent cells (1.2 μg per electroporation) (Lucigen), recovered for 1 h at 37 °C, plated across 20x 15-cm LB-Carbenicillin plates (Teknova), and incubated at 37 °C for 16 h. The total number of bacterial colonies per sub-pool was quantified using serial dilution plates to ensure a library representation of > 10,000-fold. The next morning, bacterial colonies were scraped and briefly expanded for 4 h at 37 °C in 500 ml of LB-Carbenicillin. Plasmid DNA was isolated using the Plasmid Plus Maxi Kit (Qiagen). To assess sgRNA distribution, we amplified the sgRNA target region using primers that append Illumina sequencing adapters on the 5’ and 3’ ends of the amplicon, as well as a random nucleotide stagger and unique demultiplexing barcode on the 5’ end (Table S1C). Library amplicons were size-selected on a 2.5% agarose gel, purified using the QIAquick Gel Extraction Kit (Qiagen), and sequenced on an Illumina NextSeq instrument (75 nt single end reads).

### Cell culture

Lenti-X 293T™ cells (*H. sapiens*; sex: female) obtained from Takara (cat. #632180) and Huh-7.5 cells (*H. sapiens*; sex: male) (Blight et al., 2002) were maintained at 37 °C and 5% CO2 in Dulbecco’s Modified Eagle Medium (DMEM, Fisher Scientific, cat. #11995065) supplemented with 0.1 mM nonessential amino acids (NEAA, Fisher Scientific, cat. #11140076) and 10% hyclone fetal bovine serum (FBS, HyClone Laboratories, Lot. #AUJ35777). Both cell lines have tested negative for contamination with mycoplasma.

### Lentivirus production

To produce the lentiGuidePurov2 sgRNA lentivirus library, Lenti-X 293T™ cells were seeded in 25 ml DMEM supplemented with 0.1 mM NEAA and 3% FBS at 6.25 × 10^6^ per dish in four poly-L-lysine coated p150 dishes. The following day media was removed and replaced with 20 ml fresh media and each plate was transfected with 15.6 μg lentiGuidePurov2 plasmid, 7.8 μg VSV-G plasmid, and 11.7 μg HIV Gag-Pol plasmid using Lipofectamine™ 2000 (Fisher Scientific, cat. #11668019) at 2.2 μl per ug DNA ratio supplemented with PLUS™ Reagent (Fisher Scientific, cat. #11514015) at 4.4 μl per μg DNA diluted in 6 ml Opti-MEM Reduced Serum Medium (Fisher Scientific, cat. #51985034). Six hours post transfection, the media was removed and replaced with 20 ml per plate fresh media. Twenty-four hours later media was removed and replaced with fresh media. The following day lentivirus-containing media was collected, clarified by centrifugation at 500 *g* x 5 min, filtered through a 0.22 μM filter (Corning, cat. #431153) and concentrated ∼70-fold overnight at 4 °C using Lenti-X™ concentrator (Takara, cat. #631232) following the manufacturer’s instructions. Aliquots were stored at −80 °C.

### Production and titration of coronavirus stocks

SARS-CoV-2 (strain: USA-WA1/2020) and HCoV-NL63 were obtained from BEI Resources (NR-52281 and NR-470). HCoV-OC43 was obtained from ZeptoMetrix (cat. #0810024CF) and HCoV-229E was generously provided by Volker Thiel (University of Bern). All viruses were amplified at 33 °C in Huh-7.5 cells to generate a P1 stock. To generate working stocks, Huh-7.5 cells were infected at a multiplicity of infection (MOI) of 0.01 plaque forming unit (PFU)/cell (SARS-CoV-2, HCoV-NL63, HCoV-OC43) and 0.1 PFU/cell (HCoV-229E) and incubated at 33 °C until virus-induced CPE was observed. Supernatants were subsequently harvested, clarified by centrifugation (3,000 *g* × 10 min) at 4 dpi (HCoV-229E), 6 dpi (SARS-CoV-2, HCoV-OC43) and 10 dpi (HCoV-NL63), and aliquots stored at −80 °C.

Viral titers were measured on Huh-7.5 cells by standard plaque assay. Briefly, 500 μL of serial 10-fold virus dilutions in Opti-MEM were used to infect 4 × 10^5^ cells seeded the day prior into wells of a 6-well plate. After 90 min adsorption, the virus inoculum was removed, and cells were overlaid with DMEM containing 10% FBS with 1.2% microcrystalline cellulose (Avicel). Cells were incubated for 4 days (HCoV-229E), 5 days (SARS-CoV-2, HCoV-OC43) and 6 days (HCoV-NL63) at 33 °C, followed by fixation with 7% formaldehyde and crystal violet staining for plaque enumeration. All SARS-CoV-2 experiments were performed in a biosafety level 3 laboratory.

To confirm the identity of the viruses, RNA from 200 μl of each viral stock was purified by adding 800 μl TRIzol™ Reagent (ThermoFisher Scientific, cat. #15596026) plus 200 μl chloroform then centrifuged at 12,000 *g* x 5 min. The upper aqueous phase was moved to a new tube and an equal volume of isopropanol was added. This was then added to an RNeasy mini kit column (Qiagen, cat. #74014) and further purified following the manufacturer’s instructions. Viral stocks were confirmed via next generation sequencing at the NYU Genome Technology Center using an Illumina stranded TruSeq kit and omitting the polyA selection step. Libraries were then sequenced by MiSeq Micro (2 x 250 bp paired end reads).

### Lentiviral transduction of cell lines

Huh-7.5-Cas9 cells were generated by lentiviral transduction of lentiCas9-Blast followed by selection and expansion in the presence of 5 μg/ml blasticidin. To deliver the lentiGuidePurov2 sgRNA library, 3.6 × 10^7^ Huh-7.5-Cas9 cells were transduced by spinoculation at 1,000 *g* × 1 h in media containing 4 μg/ml polybrene (Millipore, cat. #TR-1003-G) and 20 mM HEPES (Gibco, cat. #15630080) at a MOI = 0.14 to achieve ∼1,500-fold overrepresentation of each sgRNA. Cells were spinoculated at 3 × 10^6^ cells/well in 12-well plates in 1.5 ml final volume. Six hours post transduction, cells were trypsinized and transferred to T175 flasks at 6 × 10^6^ cells/flask. Two days later, media was replaced with fresh media containing 1.5 μg/ml puromycin and expanded for six days prior to seeding for coronavirus infection.

### CRISPR-Cas9 genetic screen

Huh-7.5-Cas9 cells transduced with the lentiGuidePurov2 sgRNA library were seeded in p150 plates at 4.5 × 10^6^ cells/plate in triplicate for each condition (mock, HCoV-229E, HCoV-NL63, HCoV-OC43, and SARS-CoV-2). The following day, the media was removed and viruses diluted in 10 ml/plate OptiMEM were added to cells. The inocula of HCoV-229E, HCoV-NL63 and SARS-CoV-2 were supplemented with 1 μg/ml TPCK-treated trypsin (Sigma-Aldrich, cat. #T1426) increasing the rate of infection. After two hours on a plate rocker at 37 °C, 10 ml/plate media was added and plates were moved to 5% CO2 incubators set to 33 °C or 37 °C. Coronavirus screens were performed at the following temperatures and MOIs in PFU/cell: HCoV-229E = 0.05 at 37 °C; HCoV-NL63 = 0.01 at 33 °C; HCoV-OC43 = 1 at 33 °C; SARS-CoV-2 = 0.25 at 33 °C and 37 °C. Mock cells cultured at both temperatures were passaged every 3-4 days and re-seeded at 4.5 × 10^6^ cells/plate. Media was changed on virus infected plates as needed to remove cellular debris. Mock cells and cells that survived coronavirus infection were harvested approximately two weeks post infection.

Genomic DNA (gDNA) was isolated via ammonium acetate salt precipitation if greater than 1.5 × 10^6^ cells were recovered or using the Monarch Genomic DNA Purification kit (NEB) if fewer per the manufacturer’s instructions. gDNA concentrations were quantitated via UV spectroscopy and normalized to 250 ng/μl with 10 mM Tris using a MANTIS Liquid Handler (Formulatrix). The library was amplified from gDNA by a two-stage PCR approach. For PCR1 amplification, gDNA samples were divided into 50 ul PCR reactions. Each well consisted of 25 μl of NEB Q5 High-Fidelity 2X Master Mix, 2.5 ul of 10 μM forward primer Nuc-PCR1_Nextera-Fwd Mix, 2.5 ul of 10 μM reverse primer Nuc-PCR1_Nextera-Rev Mix and 20 μl of gDNA (5 μg each reaction). PCR1 cycling settings: initial 30 s denaturation at 98 °C; then 10 s at 98 °C, 30 s at 65 °C, 30 s at 72 °C for 25 cycles; followed by 2 min extension at 72 °C. PCR1 samples were cleaned up with SPRI beads and normalized to 20 ng/μl. Each PCR2 reaction consisted of 25 μl of NEB Q5 High-Fidelity 2X Master Mix, 2.5 μl 10 μM Common_PCR2_Fwd primer, and 2.5 ul of 10 μM reverse i7 indexing primer. PCR2 cycling settings: initial 30 s at 98 °C; then 10 s at 98 °C, 30 s at 65 °C, 30 s at 72 °C for 13 cycles. PCR products were again purified by SPRI, pooled and sequenced on an Illumina NextSeq 500 at the NYU Genome Technology Center using standard Nextera sequencing primers and 75 cycles.

### Analysis of CRISPR-Cas9 genetic screen data

FASTQ files were processed and trimmed to retrieve sgRNA target sequences followed by enumeration of sgRNAs in the reference sgRNA library file using MAGeCK (Li et al., 2014). MAGeCK was also used to determine gene essentiality (beta) using its maximum likelihood estimation (MLE) algorithm. Z-scores for visualization in the form of heatmaps were computed using the following approach: for each condition, the log2 fold change with respect to the initial condition was computed. A natural cubic spline with 4 degrees of freedom was fit to each pair of infected and control cells and residuals were extracted. To obtain gene-wise data, the mean residuals for each group of sgRNAs was calculated, a z-score was computed, and a p-value was determined using a 2-sided normal distribution test. P-values were combined across screens using Fisher’s sumlog approach and corrected for multiple testing using the method of Benjamini & Hochberg.

### Analysis of RNAseq and scRNAseq data

RNAseq data from Huh-7.5 cells was obtained from GSE64677 (Luna et al., 2015) and processed using kallisto (Bray et al., 2016) followed by a tximport (Soneson et al., 2015) transcript-to-gene level transformation and then a limma-Voom approach (Ritchie et al., 2015) for read normalization. Transcript indices for kallisto were created from FASTA files from Ensembl release 100 for hg38 for all annotated cDNA and ncRNA transcripts. For Calu-3 cells, data was obtained from GSE148729 (Wyler et al., 2020). Specifically, count tables for poly-A selected libraries from bulk RNA sequenced Calu-3 cells were used and processed with the same limma-Voom read normalization scheme above. Throughout, Ensembl gene IDs were used to avoid gene symbol misidentifications. For scRNAseq analysis, Seurat objects were downloaded from FigShare (https://doi.org/10.6084/m9.figshare.12436517) (Chua et al., 2020). Using Seurat (Stuart et al., 2019), average expression of single cell data by cell type was calculated using the AverageExpression() function with default parameters. Row normalized z-scores of these average expression values were plotted per cell type for all high confidence CRISPR hits.

### Statistical Analyses

Statistical tests were used as indicated in the figure legends. Generation of plots and statistical analyses were performed using the R statistical computing environment. Error bars represent standard deviation, unless otherwise noted. We used Student’s *t*-test (unpaired, two-tailed) to assess significance between treatment and control groups, and to calculate P values. P < 0.05 was considered statistically significant.

## DATA AND CODE AVAILABILITY

Data supporting the findings of this study are reported in Supplementary Figures S1-S2 and Tables S1-S2. All raw data corresponding to high-throughput approaches (CRISPR screens, RNA-Seq) will be available through NCBI GEO. All reagents and materials generated in this study will be available to the scientific community through Addgene and/or MTAs.

---

**Supplementary Figure S1.**
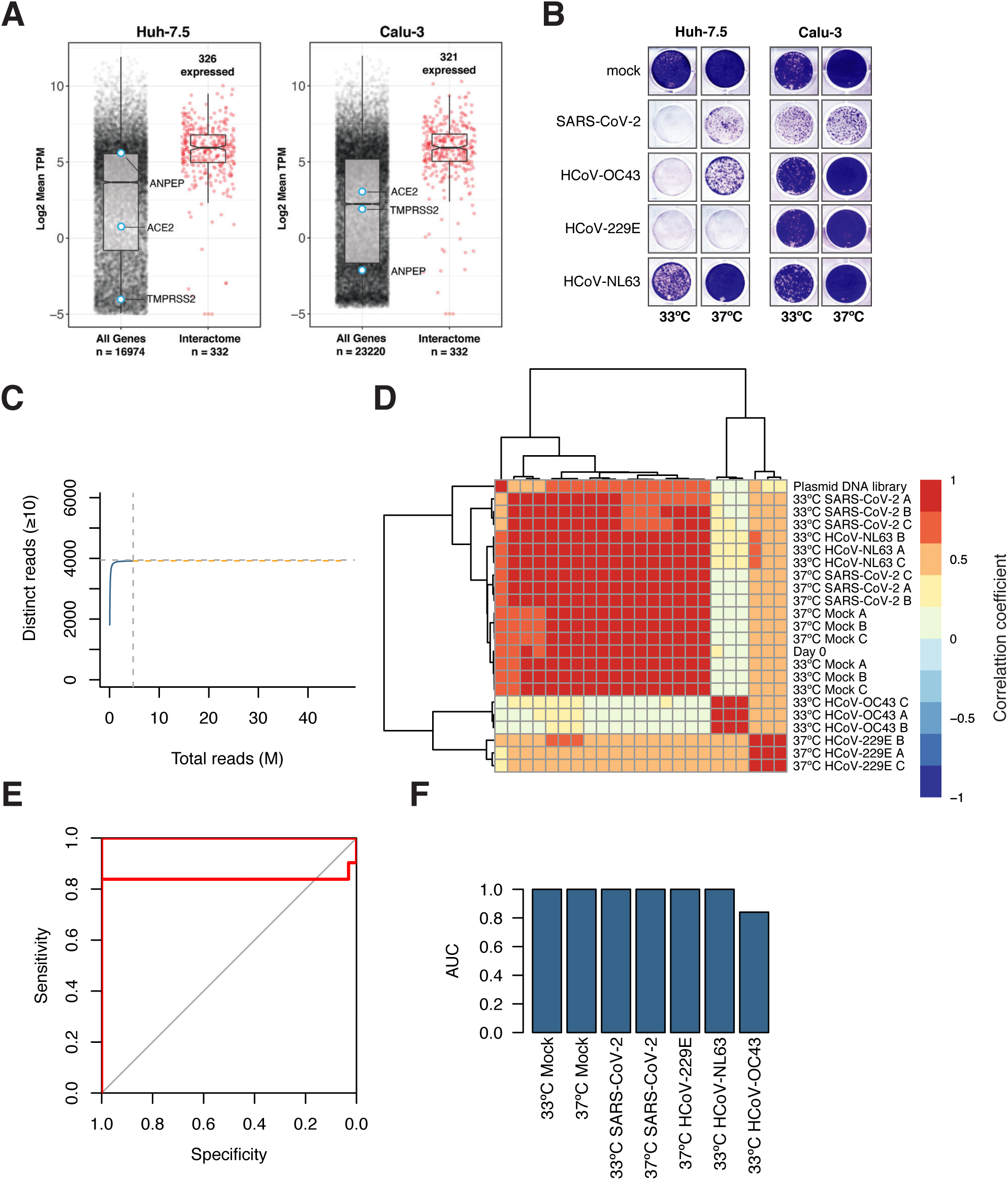
(related to Figure 1). Interactome screening cell type selection and initial quality control. **(A)** Expression profile distributions of all 332 interactome genes in Huh-7.5 or Calu-3 cells are shown in red. The expression distributions for all genes are also shown with coronavirus entry factors highlighted. **(B)** Crystal violet staining of infected cells at 7 days post infection. Huh-7.5 or Calu-3 cells were infected with SARS-CoV-2 (MOI = 0.5 PFU/cell) or endemic coronaviruses HCoV-OC43 (MOI = 1 PFU/cell), HCoV-229E (MOI = 0.5 PFU/cell), and HCoV-NL63 (MOI = 0.01 PFU/cell) at either 33 °C or 37 °C. **(C)** Species accumulation curve for each sgRNA observed more than 10 times for actual (blue) and projected sequenced reads (orange). Horizontal dashed line indicates library size. Vertical dashed line indicates number of reads sequenced. **(D)** Heatmap of Pearson correlation coefficients of normalized sgRNA counts. **(E)** Receiver operating characteristic (ROC) curves **(F)** Bar charts indicating the area under the curve (AUC) for each ROC curve in E.

**Supplementary Figure S2.**
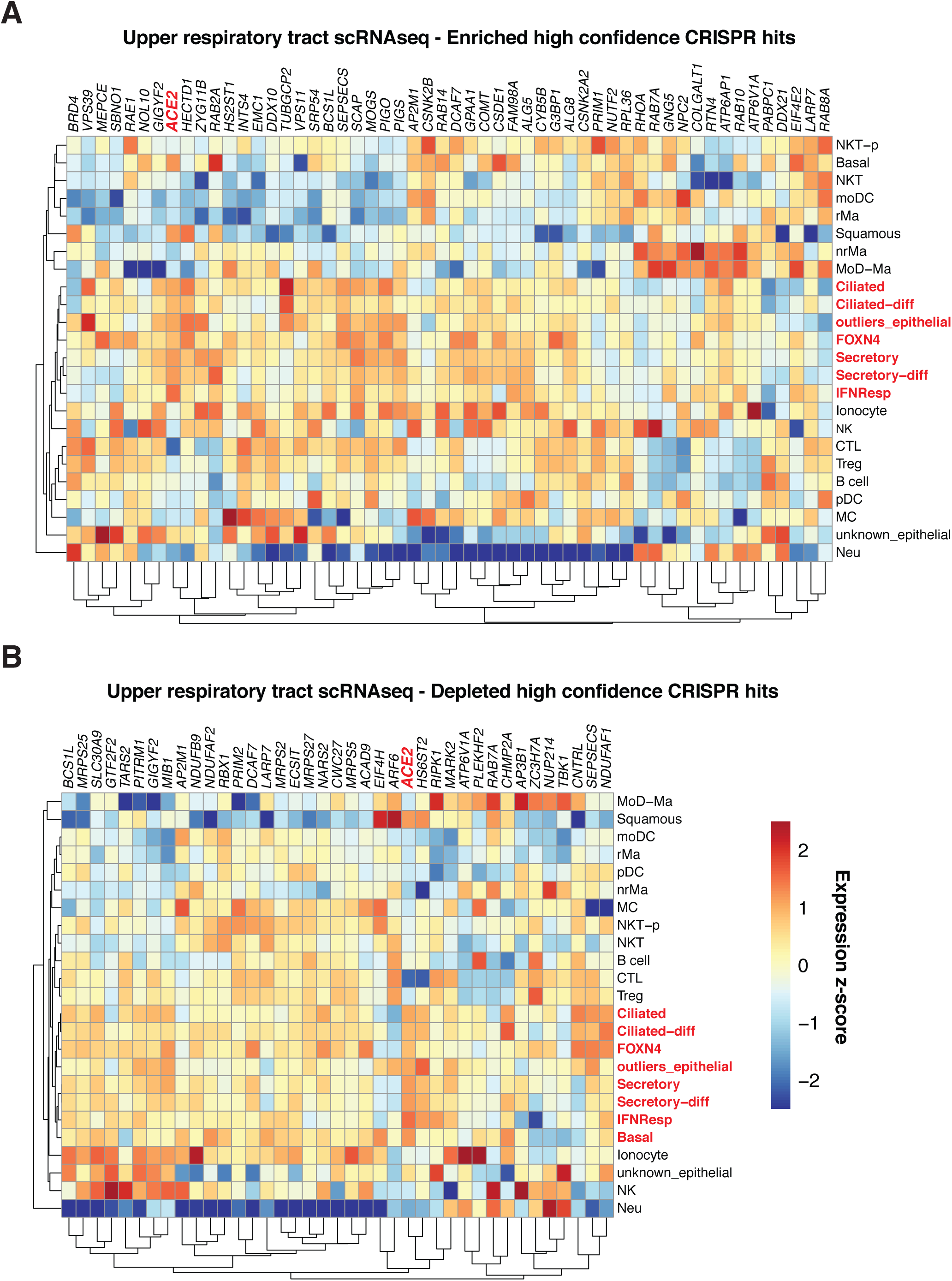
(related to Figure 4). High confidence CRISPR hits are expressed in the airway. **(A-B)** Heatmaps depicting mean expression z-scores for enriched **(A)** or depleted **(B)** CRISPR hits per indicated cell type cluster from upper respiratory tract scRNAseq in humans from (Chua et al., 2020). ACE2 and SARS-CoV-2 susceptible cell types are highlighted in red.

